# Comparative genomic analysis revealed rapid differentiation in the pathogenicity-related gene repertoires between *Pyricularia oryzae* and *Pyricularia penniseti* isolated from a *Pennisetum* grass

**DOI:** 10.1101/360016

**Authors:** Huakun Zheng, Zhenhui Zhong, Mingyue Shi, Limei Zhang, Lianyu Lin, Yonghe Hong, Tian Fang, Yangyan Zhu, Jiayuan Guo, Limin Zhang, Jie Fang, Hui Lin, Justice Norvienyeku, Xiaofeng Chen, Guodong Lu, Hongli Hu, Zonghua Wang

**Author notes:** These authors contributed to this work equally (Huakun Zheng, Zhenhui Zhong). **Corresponding author: Zonghua Wang**, Mailing address: Institute of Oceanography, Minjiang University, Fuzhou 350108, China; **Hongli Hu**; **Guodong Lu**, National Engineering Research Center of JUNCAO Technology, College of Life Science, Fujian Agriculture and Forestry University, Fuzhou 350002, China. State Key Laboratory of Ecological Pest Control for Fujian and Taiwan Crops, College of Plant Protection, Fujian Agriculture and Forestry University, Fuzhou 350002, China.

## Abstract

**Backgrounds:** *Pyricularia* is a multispecies complex that could infect and cause severe blast disease on diverse hosts, including rice, wheat and many other grasses. Although the genome size of this fungal complex is small [~40 Mbp for *Pyricularia oryzae* (syn. *Magnaporthe oryzae*), and ~45 Mbp for *P. grisea*], the genome plasticity allows the fungus to jump and adapt to new hosts. Therefore, deciphering the genome basis of individual species could facilitate the evolutionary and genetic study of this fungus. However, except for the *P. oryzae* subgroup, many other species isolated from diverse hosts, such as the *Pennisetum* grasses, remain largely uncovered genetically.

**Results:** Here, we report the genome sequence of a pyriform-shaped fungal strain *P. penniseti* P1609 isolated from a *Pennisetum* grass (JUJUNCAO) using PacBio SMRT sequencing technology. We performed a phylogenomic analysis of 28 Magnaporthales species and 5 non-Magnaporthales species and addressed P1609 into a *Pyricularia* subclade that is distant from *P. oryzae*. Comparative genomic analysis revealed that the pathogenicity-related gene repertoires were fairly different between P1609 and the *P. oryzae* strain 70-15, including the cloned avirulence genes, other putative secreted proteins, as well as some other predicted *Pathogen-Host Interaction* (*PHI*) genes. Genomic sequence comparison also identified many genomic rearrangements.

**Conclusion:** Taken together, our results suggested that the genomic sequence of the *P. penniseti* P1609 could be a useful resource for the genetic study of the *Pennisetum*-infecting *Pyricularia* species.

## Introduction

*Pyricularia* was established by Saccardo to accommodate a type of fungal species based on the pyriform conidia when the first species of this pathogen, *Pyricularia grisea*, was isolated from crabgrass (*Digitaria sanguinalis* L.) [1]. What raised the concern of this *Pyricularia* fungus was the notorious blast disease on rice and wheat caused by one of its species, *Pyricularia oryzae* (syn. *Magnaporthe oryzae*) [2-4]. To date, 100 plant genera comprising of 256 species have been documented as the hosts of the *Pyricularia* species (https://nt.ars-grin.gov/fungaldatabases/fungushost/fungushost.cfm), among which 54 genera belong to the Poaceae family. Seven *Pyricularia* species (including one unidentified species) have been isolated from *Pennisetum* spp., a widespread genus in the Poaceae family, and more than one *Pyricularia* species can be found on the same *Pennisetum* species. For instance, 4 *Pyricularia* species, namely, *P. penniseti*, *P. penniseticola*, *P. setariae* and *Pyricularia sp*. have been found on *P. typhoides* [5, 6].

The genome sequence of *P. oryzae* strain 70-15, a hybrid clone of rice-infecting isolate 104-3 and the weeping love grass isolate AR4 [7] has facilitated the revelation of developmental and pathogenic mechanisms of the blast fungus and rendered it to become one of the most important model fungus [8, 9]. Since the publication of the *P. oryzae* strain 70-15 genome, more field blast isolates were sequenced and assembled, including the Ina168, HN9311, FJ81278, Y34, P131, 98-06 and Guy11 [10-14]. Comparative genomic analyses and functional studies of these strains revealed genome plasticity and the involvement of the lineage specific genes in pathogenicity [12, 14]. More recently, facilitated by the fast developing of sequencing technologies, a number of field isolates from rice, as well as isolates from different grass and cereal hosts, were sequenced and subjected to population-level analyses, revealing host immunity as the major force driving specialization after host shift [2, 15-18]. However, genomes of most of the species of the *Pyricularia* complex remain unexplored. For example, among the 7 identified *Pyricularia* species isolated from *Pennisetum* grasses [5], only *P. pennisetigena* was recently sequenced [2].

Here, we reported the whole-genome sequence of *P. penniseti* [19] isolated from a *Pennisetum* grass JUJUNCAO (*Pennisetum giganteum* z. x. Lin). JUJUNCAO was originally developed as culture matrix for the cultivation of edible mushrooms by Lin *et al*, and later became a versatile grass that are used as forage for cattle and sheep, material for the biofuel production, and tool for the remedy of soil erosion [20-23]. We have recently isolated a pyriform-shaped fungus, P1609, from leaf spots of JUJUNCAO. P1609 caused a typical blast fungal disease symptom on JUJUNCAO, showing small, round or elliptical lesions as an initial symptom and spindle shaped, grayish to tan necrotic with yellow halos at a later disease stage. Since the morphologic and phylogenetic analyses distinguished P1609 from other identified *Pyricularia* species, but undistinguished from the *P. penniseti* reported in 1970 India [5, 24], we therefore termed this fungus *P. penniseti* [19]. In this study, we performed genome sequencing of this strain, aiming for a proper classification of this fungus in the *Pyricularia* population and identification of genes that may be involved in the adaptation of this fungus to JUJUNCAO.

## Results

### Genome sequencing and assembly

We sequenced the P1609 genome with the long-read PacBio technology. In total, 312,061 reads with 8.6 Kb average lengths were obtained, representing about 60-fold coverage of the genome (Fig. 1A). The genome sequence was assembled with the HAGP pipeline, resulting in the total assembly space of 41.82 Mb (Table 1), similar to assemblies of isolates sequenced by PacBio [10]. The assembly contains 53 contigs, with the N50 of 3.4 Mb and the largest contig of 7.56 Mb (Fig. 1B). Contigs > 1Mb cover 89.7% and contigs > 100 Kb cover 98.5% of the genome (Fig. 1B; Table 1), indicating long-continuity of the assembly. The GC content of the assembly is 50.3%, similar to genomes of *Pyricularia* isolates from different host plants, which range from 48.6% to 51% [18]. Genome annotation identified 13,102 genes with average gene size of 1,758 bp, fairly evenly dispersed on contigs (Table 2, Fig. 1C, track b).

**Fig. 1.**
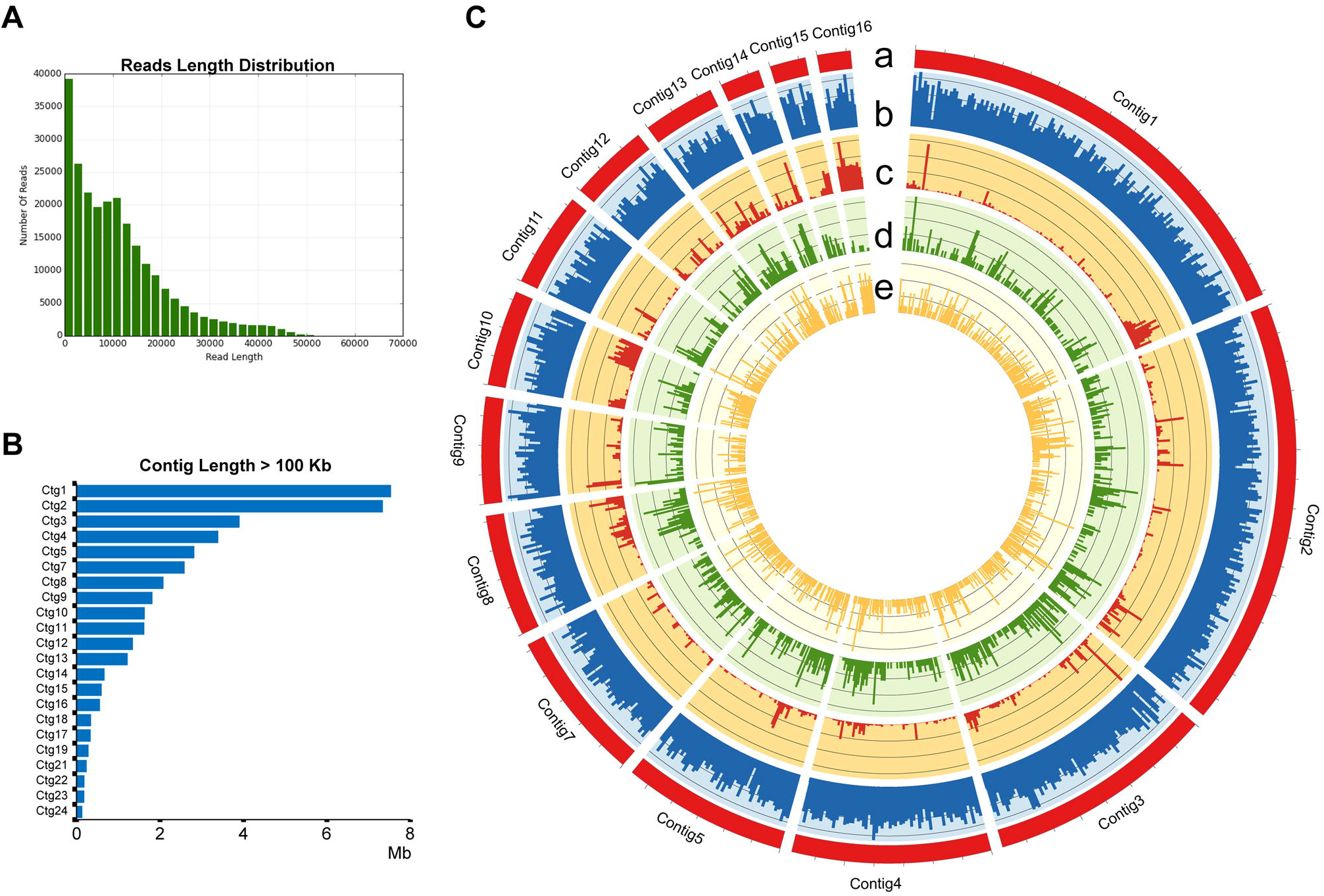
PacBio sequencing and genome assembly of P1609. (A) Reads length distribution. (B) Contig length of assembled contig > 100 Kb. (C) Overview of P1609 genome. (Track a) Contig1 to contig16 of P1609, (track b) gene density, (track c) Transposon elements density, (track d) secreted proteins density, (track e) unique gene (compared with *P. oryzae*, *N. crassa*, *F. graminearum* and *C. gloeosporioides*) density of P1609 per 50 Kb.

**Table 1:** Details of sequencing reads and genome assembly of P1609

| **Feature** | **Value** |
| --- | --- |
| Number of Subreads | 312,061 |
| Total Length of Subreads (bp) | 2,688,966,115 |
| Mean Coverage | 59 |
| Polished Contigs | 53 |
| Maximum Contig Length (bp) | 7,555,856 |
| N50 Contig Length (bp) | 3,411,241 |
| Sum of Contig Lengths (Mb) | 41.82 |
| GC level | 50.30% |
| length > 1 Mb | 89.70% |
| length > 100 Kb | 98.50% |

**Table 2:** Details of genome annotation of P1609

| Category | Value |
| --- | --- |
| Total TE | 7.67% |
| LINEs | 1.50% |
| LTR elements | 4.27% |
| DNA elements | 0.63% |
| Unclassified | 1.27% |
| Gene Number | 13102 |
| Average Gene Length (bp) | 1758 |
| Secreted Protein Number | 1409 |

*De novo* repeat sequence analysis identified 7.67% of repeat sequences, among which 4.27% were Gypsy and Copia, the two long terminal repeats (LTR)-type retrotransposons. This finding is consistent with former studies, suggesting that LTR-type retrotransposons are the most expanded transposable elements (TEs) in P1609 genome [25]. The TEs are not evenly distributed along the contigs. While some contigs are enriched with TEs, other contigs have less TE rich regions, suggesting that the P1609 genome also underwent transposon expansion as observed in other plant pathogens [26].

### Comparative and phylogenetic analysis

To understand the genetic relationship of P1609 with other fungal phytopathogens, we generated a phylogenetic tree of P1609 with *Botrytis cinerea* (Bc), *Colletotrichum gloeosporioides* (Cg), *Fusarium graminearum* (Fg), *Neurospora crassa* (Nc), *Pyricularia oryzae* (Po), *Sclerotinia sclerotiorum* (Ss), *Trichoderma reesei* (Tr) and *Ustilago maydis* (Um). Since P1609 showed a close morphological relationship with *P. oryzae* isolates, we included *Pyricularia* isolates collected from different host plants, including *Oryza sativa* (PoOs), *Triticum aestivum* (PoTa), *Digitaria sanguinalis* (PoDs), *Setaria viridis* (PoSv), and *Eleusine indica* (MoEi). In total, 2,051 single-copy genes shared by all the examined genomes were selected to infer phylogeny [27]. The resulting phylogenetic tree indicated that P1609 is more divergent with PoOs, PoTa, PoSv, and PoEi (*P. oryzae*) in contrast to PoDs (*P. grisea*) (Fig. 2A). We then estimated divergence time of P1609 and *Pyricularia* isolates by assuming a constant molecular clock calibrated in a previous study, which estimated the divergence of *Neurospora* and *Pyricularia* at about 200 million years ago (MYA). The estimation indicated that P1609 and *Pyricularia* isolates diverged at about 31 MYA, earlier than the divergent time of rice- and *S. viridis*-infecting isolates (about 10,000 years ago) [28, 29]. The phylogenetic tree also indicated that P1609 is a member of Magnaporthales, but is distantly related to the *Pyriculaira* isolates collected from PoOs, PoTa, PoDs, PoSv, and PoEi. To further determine exactly where P1609 localized in Magnaporthales, we also generated a phylogenomic tree of P1609 with 28 Magnaporthales species and 5 non-Magnaporthales species using amino acid sequences of 226 conserved orthologous genes as described [30]. The result showed that P1609 was localized in the *Pyricularia* subclade, and was closer to *P. oryzae* than *Xenopyricularia zizaniicola* (Fig. 2B), indicated that P1609 is a *Pyricularia* species.

**Fig. 2.**
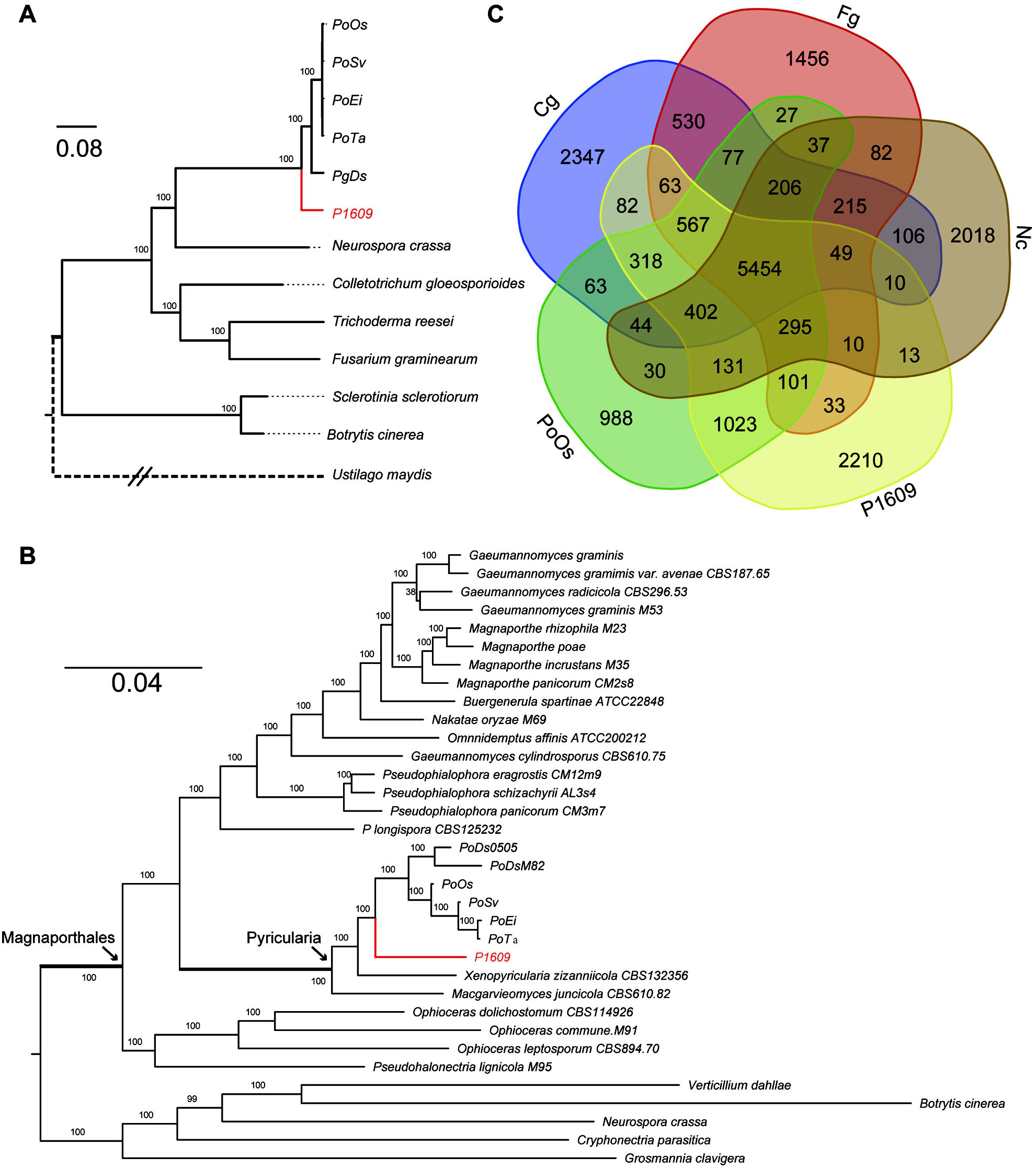
Phylogenetic and comparative genomic study of P1609. (A) Phylogenomic tree of P1609 with *Botrytis cinereal* (Bc), *Colletotrichum gloeosporioides* (Cg), *Fusarium graminearum* (Fg), *Neurospora crassa* (Nc), *Sclerotinia sclerotiorum* (Ss), *Trichoderma reesei* (Tr) and *Ustilago maydis* (Um) as well as *Pyricularia* isolates collected from *O. sativa* (PoOs), *T. aestivum* (PoTa), *D. sanguinalis* (PoDs), *S. viridis* (PoSv), and *E. indica* (PoEi) based on 2,051 single copy genes. The values of all of the branches are 100. (B) Maximum likelihood tree of P1609 and 28 Magnaporthales species, as well as 5 Sordariomycetes used as outgroup species based on 82,715 amino acid positions derived from 226 genes. (C) Venn diagram showed an overlap of gene families among P1609, *Pyricularia* rice isolates (PoOs), *C. gloeosporioides* (Cg), *F. graminearum* (Fg) and *N. crassa* (Nc).

We next conducted a comparative genomic study of P1609 with 70-15 (the reference isolate of *P. oryzae*), *N. crassa*, *F. graminearum* and *C. gloeosporioides*. Venn diagram showed that 5 organisms share 5,454 of gene families with each other, which covers 50.2% of the gene set of P1609 (Fig. 2C). Notably, the number of unique genes (no homologs in the selected organisms) in P1609 was 2,210, which is about twice more than the number of unique genes recorded for 70-15. Although most of these unique genes had no functional annotations, Pfam annotation indicated that some of the unique genes encode carbohydrate-active enzymes involved in polysaccharides metabolism pathways. The high representation of carbohydrate-active enzymes may be related to the adaptation to the host *P. giganteum* (Fig. S1). We therefore scanned natural selection of 5,991 pairs of orthologs of *P. oryzae* and P1609, identifying 6 genes with Ka/Ks > 1 (Table 3). These genes are involves in different secondary metabolic pathways.

**Table 3:** Positively selected genes in P1609

| Protein ID | Ka | Ks | Ka/Ks | P-Value (Fisher) | Description |
| --- | --- | --- | --- | --- | --- |
| P1609_7184 | 0.707979 | 0.440748 | 1.60631 | 0.00697146 | Unknown |
| P1609_5032 | 0.550905 | 0.343609 | 1.60329 | 1.98443E-06 | Isoamyl alcohol oxidase |
| P1609_3360 | 0.183224 | 0.118517 | 1.54597 | 0.00293936 | Glycerol uptake protein 1 |
| P1609_791 | 0.302231 | 0.199786 | 1.51277 | 9.75093E-06 | Folypolyglutamate synthase |
| P1609_7869 | 0.491873 | 0.328109 | 1.49911 | 0.0296442 | Thioredoxin reductase |
| P1609_1006 | 0.40206 | 0.275456 | 1.45962 | 0.0288585 | Spore surface glycoprotein BclB |

### Gene Categories Involved in Pathogenicity

Plant pathogenic fungi employed diverse gene repertoires to invade host plants and subvert host immune systems, which include effectors, carbohydrate-active enzymes (CAZymes), other secreted enzymes and fungal secondary metabolisms [31, 32]. P1609 and *P. oryzae* isolates are divergent and no cross-infectivity exists between them. To understand their secretomic difference, we first compared predicted secretomes between P1609 and the *P. oryzae* isolates. Among 1,409 putative secreted proteins of P1609, 236 were unique in P1609 (Table S1). By contrast, 165 putative effectors in the 70-15 genome were absent in the P1609 genome (Table S2). Notably, all known avirulence genes (*AVRs*) from the rice-infecting isolate genomes were absent in P1609 genome.

We next compared CAZymes of P1609, groups of enzymes enable plant pathogens to break down the plant cell wall [33]. Our BLASTp search results showed that P1609 contains more predicted CAZyme-coding genes than 70-15 (Fig. S2). Detailed analysis showed that the P1609 genome encodes 5 unique CAZyme-coding genes, namely CBM61, GH117, GH35, GH65 and PL24. While it has 6 copies of GH28 pectinases (3 copies in *Pyricularia* species; Fig. S2). We then further analyzed the distribution of annotated PHI genes in P1609. In total, we identified 1,692 potential PHI genes belonging to 1,154 gene families (Table S3). Interestingly, we found that several PHI genes exhibited great expansion in P1609 genome. For instance, *MGG_12656*, a gene involved in virulence in *P. oryzae*, has 107 homologs in P1609, and ChLae1 contributing to toxin production and virulence in a maize pathogen *Cochliobolus heterostrophus* has 17 homologs in P1609 [34, 35]. Compared with 70-15, 35 PHI genes were unique in P1609 (Table 4), most of which had highly similar homologs in *Fusarium* (*Gibberella*).

**Table 4:**
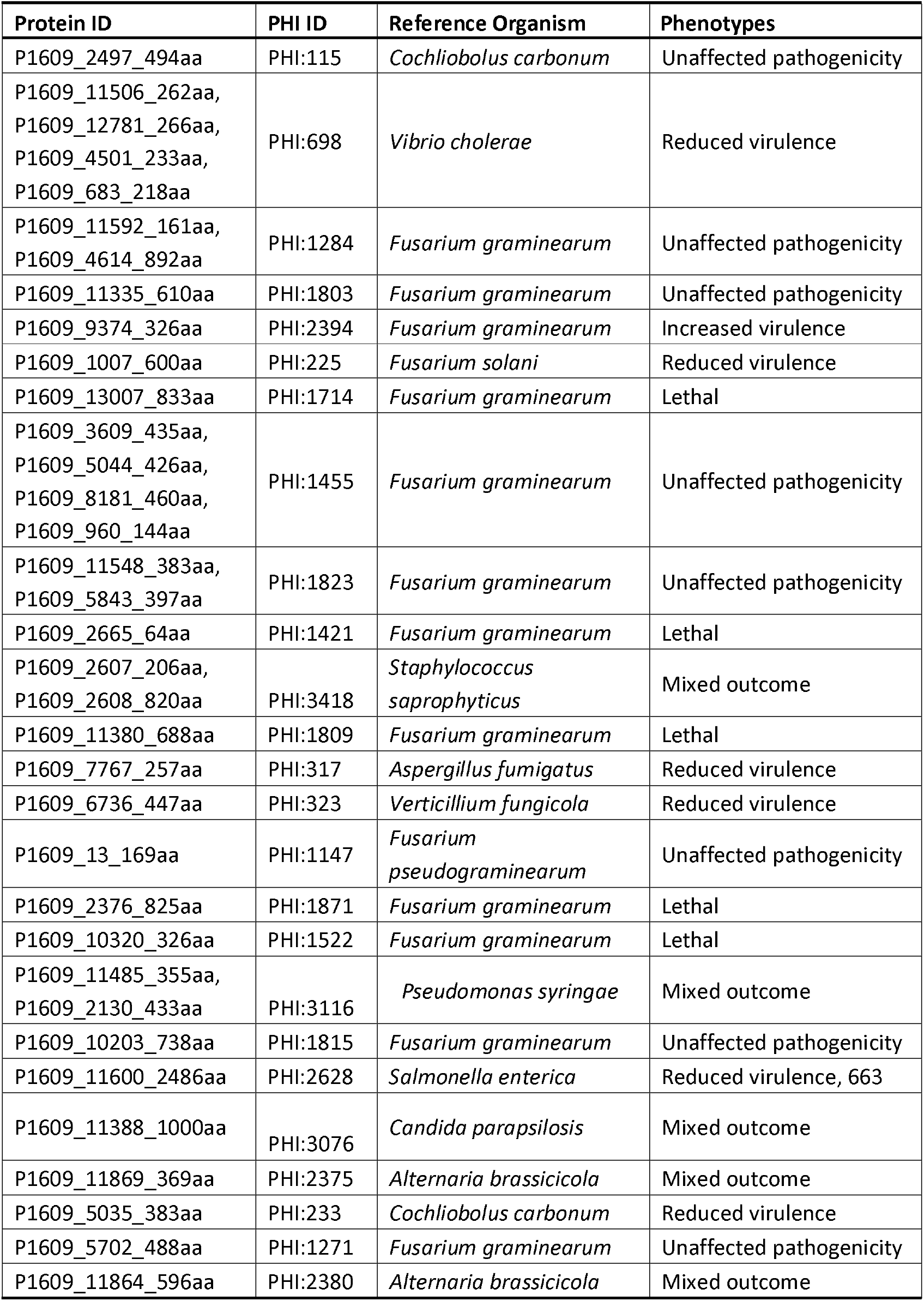
Unique Pathogen Host Interaction (PHI) genes in P1609.

### Chromosome rearrangements

To explore genome collinearity and rearrangement between P1609 and 70-15, since 70-15 was assembled into chromosome level and showed a closer phylogenetic relationship with the P1609. The identified collinear gene blocks that linked with different chromosomes in 70-15 (Fig. 3A) were visualized in the contigs (ctgs) of P1609 (Fig. 3B). Generally, P1609 and 70-15 displayed high genome collinearity. Ctg2, ctg3, ctg5 and ctg7 in P1609 correspond to chr.2, chr.6, chr.3 and chr.1 of 70-15, respectively. We found that the second largest contig in the P1609, ctg2, is a recombinant of chr.4 and chr.7 of 70-15 (Fig. 3B). The joining region was spanned by multiple single PacBio long reads (Fig. 4), excluding the possibility that the rearrangement was an artifact due to assembly errors. Meanwhile, notably, contig end region of P1609 showed a higher level of chromosome deletion and rearrangement (Fig. 3c). For instance, ctg4 and ctg8 were merged by blocks from chr.1 and chr.3 at the end of the contig, while ctg9 was merged by blocks from chr.3 and chr.5 at the end of the contig (Fig. 3B).

**Fig. 3.**
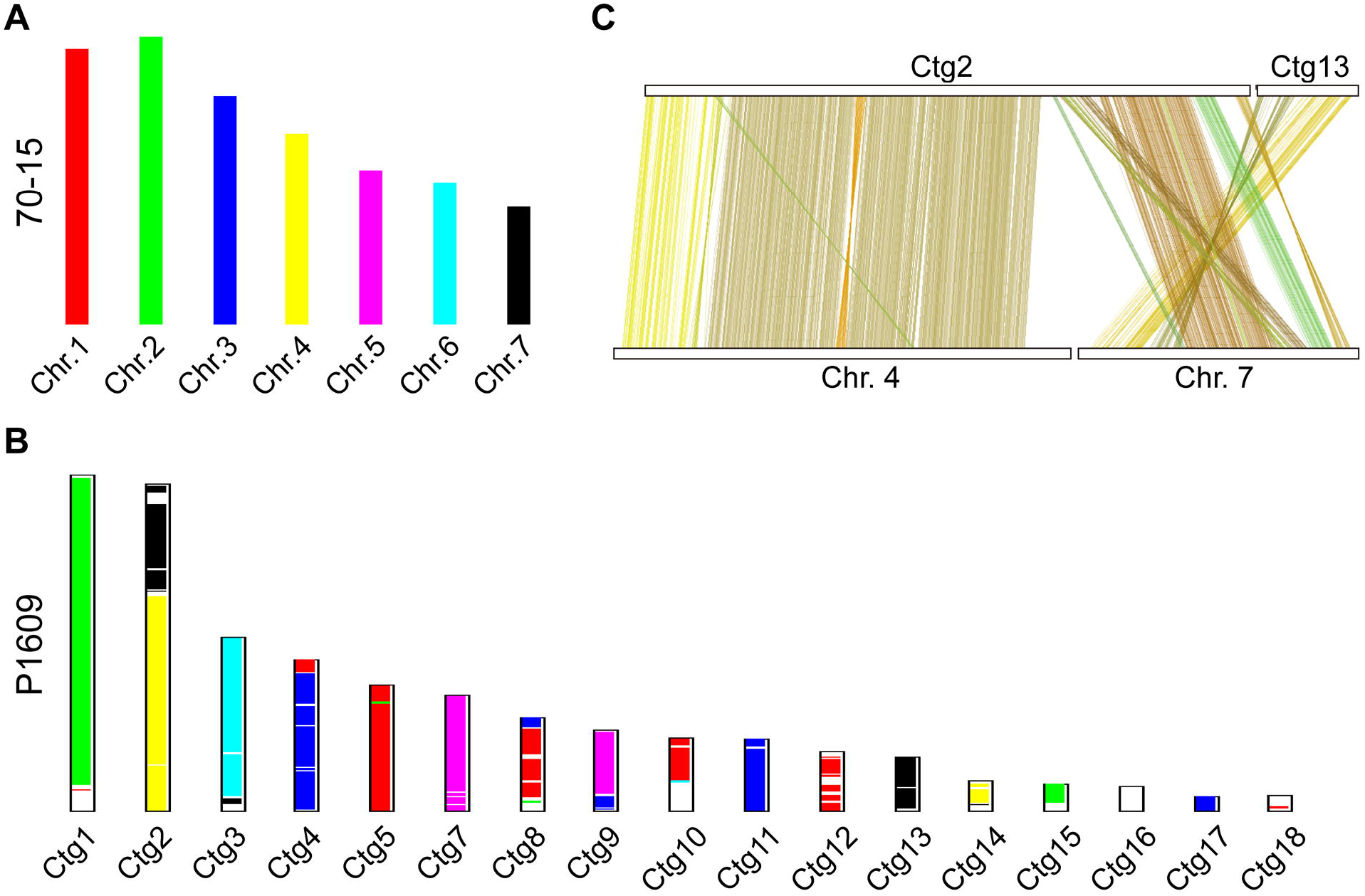
Chromosome rearrangement and splitting between P1609 and 70-15. (A) Bar plot showing the chromosomes in 70-15. (B) Bar plot showing the assembled contigs of P1609. Colinear chromosomes of 70-15 and contigs of P1609 are indicated by the same color. (C) Dual synteny plot showing splitting of Ctg2 of P1609 into chr. 4 and chr. 7 in 70-15.

**Fig. 4.**
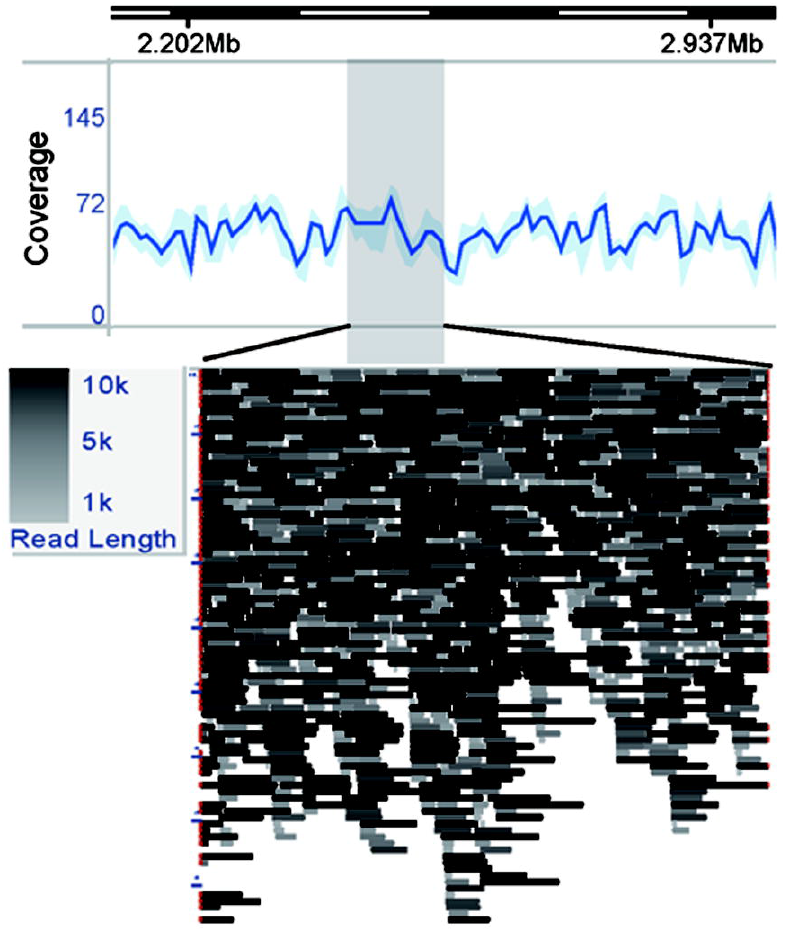
PacBio long-read coverage from 2.02 to 2.94 Mb of Ctg 2. Color of reads indicate different read lengths.

## Discussion

Although the genome size of *Pyricularia* species is small, it is well-known for its complicated genomic plasticity. Here we sequence the genome of *P. penniseti* using the PacBio SMRT technology. Comparative genomic analysis revealed several chromosome rearrangement events and difference in pathogenicity-related gene repertoires between the *P. penniseti* strain P1609 and the *P. oryzae* strain 70-15.

## Chromosome fission and fusion

Long-read sequencing greatly improved genome assembly and thus provided more valuable details on genome structures at a chromosome level. By comparing the chromosome structure between P1609 and 70-15, we found several chromosome splitting and rearrangement events. Chromosomal rearrangements have been reported to be associated with virulence evolution in several pathogens by losing the *AVR* gene(s) [26, 36, 37]. Our results indicated frequent chromosome rearrangement and splitting in the blast fungi, and the telomere regions are very unstable in its haploid genome during the adaptation to different host species. Further study is required to investigate the role of the chromosome recombination during the adaptation to JUJUNCAO.

## Unique genes and positively selected genes

In this study, we found 2,210 unique genes that were absent in the 70-15 genome. The P1609 genome encodes more CAZymes than 70-15 genome. For example, P1609 contains twice the number of genes encoding glycosyl hydrolase family 28 (GH28) pectinases in 70-15. Generally, necrotrophic plant pathogens possess more GH28 enzymes than biotrophic and non-pathogens fungi [38]. These results suggested that P1609 might heavily rely on CAZymes in the interaction with JUJUNCAO.

To identify the positively selected genes, we detected with Ka>Ks from 5,991 orthologous pairs between *P. oryzae* and P1609 and identified 6 genes with Ka>Ks in P1609. Functional annotation showed that most of these genes involved in secondary metabolic pathways. For example, P1609_5032 encodes an isoamyl alcohol oxidase that turns isoamyl alcohol into isovaleraldehyde [39]. In *Saccharomyces cerevisiae*, isoamyl alcohol could induce filament formation [40]. P1609_5032 might play a role in fungal development through controlling the level of isoamyl alcohol. P1609_791 encodes a folylpolyglutamate synthase involved in biosynthesis of folate, required for protein synthesis of bacterial, mitochondrial, and chloroplast, including purines, dTMP, methionine, and formyl-methionyl-tRNA [41]. P1609_3360 encodes a glycerol uptake protein, which is an important intracellular osmolyte participating in osmotic stress response [42] and critical for the function of appressorium in *P. oryzae* [43]. P1609_7869 encodes a spore surface glycoprotein. Its homologs have been proved to be involved in spore adhesion to hydrophobic surface in several *Colletotrichum* species [44-46] rather than spore tip mucilage as in *P. oryzae* [47]. P1609_1006 encodes a BclB glycoprotein (collagen-like protein). Its homologs in *Bacillus anthracis* were important components of the infection-associated structure exosporium [48-50], although its role in filamentous pathogens remains unknown. The positive selection on these genes suggested that they might play roles in the interaction between P1609 and JUJUNCAO.

## Putative secreted proteins

There are two layers of plant immunity: the pathogen-associated molecular patterns (PAMPs)-triggered immunity (PTI), and the effetor-triggered immunity (ETI). It was previously proposed by Schulze-Lefert and Panstruga that ETI is the major force driving the host specificity of pathogens [51]. Our previously study focusing on *AVR* gene evolution of the *Pyricularia* species also revealed that directional selection exerted by host plants is the direct force driving host specificity in *Pyricularia* species [18]. Recent studies on the *P. oryzae* populations revealed that divergent host immunity systems (both PTI and ETI) in japonica (*Oryza sativa* subsp. xian) and indica (*Oryza sativa* subsp. geng), determining the deposition of effector repertoires and specialization to the two subspecies [15-17]. Our comparative genomic analysis showed that P1609 contains a large number of unique effector candidates by comparing with 70-15, but lost many putative effectors in the 70-15 genome, including all known AVR effectors. One possible explanation is that the JUJUNCAO harbors a high level of basal immunity, as well as an arsenal of resistance genes, which driven P1609 to gain a lot of effectors to overcome the robust basal immunity posed by JUJUNCAO, and at the meanwhile, abandoned the *AVR* genes that could be recognized by the *R* genes from JUJUNCAO.

## Conclusion

*Pyricularia* species are pathogens of either food- or forage grasses. The model fungus *P. oryzae* had been well studied. However, there are only a few whole-genome sequences available of other species, such as those from *Pennisetum* grasses. Here, we generated long-read PacBio reads and produced a assemblage with long-continuity contig sequences. Phylogenomic and comparative genomic analysis showed that P1609 is a *Pyricularia* species genetically distant with *P. oryzae*, and the two genomes vary substantially in their pathogenicity-related gene repertoires. In summary, the P1609 assembly and genome annotation represents the few available *Pyricularia* genome resources for studying the pathogenic mechanism of this fungus towards *Pennisetum* grasses.

## MATERIALS AND METHODS

### Isolation of the fungal strain

The *Pennisetum*-infecting strain P1609 was isolated from the leaf spot lesion of JUJUNCAO (*Pennisetum giganteum* z. x. Lin), in the nursery of National Engineering Research Center of JUNCAO Technology, Fujian Agriculture and Forestry University located at No. 63 Xiyuangong Road, Minhou County, Fuzhou, Fujian Province, China.

### DNA extraction, amplification and sequencing

To prepare the genomic DNA for sequencing, the P1609 isolate was cultured in the liquid complete medium (CM) in a 110-rpm shaker at 25 °C for 3 to 4 days. The mycelia were then collected for the preparation of genomic DNA using a CTAB method as previously described [18]. Sequencing libraries were prepared using the SMRTbellTM Template Prep Kit 1.0 (PACBIO) and sequenced using PacBio Sequel platform (NovoGene, China).

### Assembly and annotation

*De novo* sequence assembly was conducted by SMRTLink v. 5.0.1.10424, HGAP 4 pipeline provided by Pacific Bioscience Company. In HGAP 4 pipeline, the expected genome size was set as 45 Mb based on the reported size of *Pyricularia* genomes, and default settings were used for other parameters. Gene prediction was conducted using Fgenesh from SoftBerry (MolQuest II v2.4.5.1135, http://linux1.softberry.com/berry.phtml) with *Pyricularia* additional variants as training organism. Gene functional domain annotation was conducted by InterproScan (version 4.8, http://www.ebi.ac.uk/interpro/), and PfamScan [52]. Pathogen-Host Interaction (PHI) genes were predicted by performing a whole genome blastp analysis against the PHI database (E<10^−10^) [53, 54]. Putative carbohydrate-active enzymes (CAZymes) were identified using the HMMER 3.1b1 by searching annotated HMM profiles of CAZymes downloaded from the dbCAN database in protein sequences of P1609 [55].

### Repeat analysis

*De novo* repeat sequence identification was analyzed by using RepeatModeler (version 1.0.8) with default settings. Repeat sequences obtained from RepeatModeler have been used to search for repeat sequences in the P1609 genome by RepeatMasker (version 3.3.0) (http://www.repeatmasker.org/) [56].

### Phylogenetic analysis and comparative genomic analysis

Phylogenomic tree of P1609 and *B. cinerea* [57], *C. gloeosporioides* [58], *F. graminearum* [59], *N. crassa* [60], *P. oryzae* [8], *S. sclerotiorum* [57], *T. reesei* [61] and *U. maydis* [62] was built based on single copy orthologs from clustering result of OrthoFinder (v0.6.1) [27]. 2,051 single copy genes have been selected out from 13 organisms in total (see Fig. 1A) and aligned with MAFFT (mafft-linsi-anysymbol) [63]. The phylogenetic tree was constructed using FastTree based on the alignments of single-copy orthologs with approximately-maximum-likelihood model and bootstrap 100 [64]. For divergence time estimation, the phylogenetic analysis was conducted using r8s (version 1.81), and the divergence time of *Pyricularia* and *Neurospora* (200 MYA) was used as a reference [29, 65]. Clustering result of 13 genomes was also used for unique gene identification and comparative genomic analysis. We used MCScanX to identified syntenic blocks between P1609 and 70-15. To detect the conserved synteny blocks, the reciprocal best-match paralogs of P1609 and 70-15 were conducted by all-against-all BLASTP comparison, with E-value <10^−10^ [66].

### List of abbreviations

**Bc:** *Botrytis cinerea* (*B. cinerea*); **Cg:** *Colletotrichum gloeosporioides* (*C. gloeosporioides*); **Fg:** *Fusarium graminearum* (*F. graminearum*); **Nc:** *Neurospora crassa* (*N. crassa*); ***P. grisea:*** Pyricularia grisea; ***Po:*** Pyricularia oryzae (*P. oryzae*); **Ss:** *Sclerotinia sclerotiorum* (*S. sclerotiorum*); ***Tr:** Trichoderma reesei* (*T. reesei*); **Um:** *Ustilago maydis* (*U. maydis*); **PoDs:** *Pyricularia* strain isolated from *Digitaria sanguinalis*; **PoSv:** *Pyricularia* strain isolated from *Setaria viridis*; **MoEi:** *Pyricularia* strain isolated from *Eleusine indica*; **PoOs:** *Pyricularia* strain isolated from *Oryza sativa*; **PoTa:** *Pyricularia* strain isolated from *Triticum aestivum*; ***P. penniseti:** Pyrucularia penniseti*; ***P. giganteum:** Pennisetum giganteum*; ***P. penniseticola:** Pyricularia penniseticola*; ***P. setariae:** Pyricularia setariae*;***P. typhoides***: *Pennisetum typhoides*;***AVR gene:*** avirulence gene; **CAZymes:** carbohydrate-active enzymes; **CBM:** Carbohydrate-Binding Module Family; chr: chromosome; **CM:** complete medium; **CTAB:** Cetyltrimethyl Ammonium Bromide; **ctg:** contig; **ETI:** effetor-triggered immunity; **GH:** glycosyl hydrolase family pectinases; **LTR:** long terminal repeats; **MYA:** million years ago; **PHI:** Pathogen-Host Interaction; **PL:** Polysaccharide Lyase Family; **PTI:** pathogen-associated molecular pattern (PAMPs)-triggered immunity; ***R genes:*** resistance gene; **SMRT:** Single-Molecule Real-Time; **TE:** transposable elements.

## Declarations

### Ethics approval and consent to participate

Not applicable.

### Consent for publication

Not applicable.

### Availability of data and materials

Genome assembly and PacBio reads are available in GenBank under BioProject PRJNA416656. The Whole Genome sequence has been deposited at GenBank under the accession PELF00000000.

## Authors’ contributions

The study was conceived and designed by ZW and GL. The initial collection and culturing of the strain was performed by HZ, XC, HH, MS, LZ, TF, YZ, JG, LZ, JF and HL. Bioinformatics was performed by ZZ, and LL. ZZ, HZ, JN, GL and ZW wrote, revised and approved the manuscript. All authors read and approved the final manuscript.

## Acknowledgments

We would like to thank Dr. Sanzhen Liu at Kansas State University for critically reading the manuscript.

## Open Access

This article is distributed under the terms of the Creative Commons Attribution 4.0 International License (http://creativecommons.org/licenses/by/4.0/), which permits unrestricted use, distribution, and reproduction in any medium, provided you give appropriate credit to the original author(s) and the source, provide a link to the Creative Commons license, and indicate if changes were made. The Creative Commons Public Domain Dedication waiver (http://creativecommons.org/publicdomain/zero/1.0/) applies to the data made available in this article, unless otherwise stated.

## Additional files

**Additional files 1: Table S1.** P1609_vs_7015_unique_secreted

**Additional files 2: Table S2.** 70-15 VS P1609 unique secreted proteins

**Additional files 3: Table S3.** Predicted PHI in P1609

**Additional files 4: Figure S1.** Gene copy numbers of CAZYmes in *Botrytis cinereal* (Bc), *Colletotrichum gloeosporioides* (Cg), *Fusarium graminearum* (Fg), *Neurospora crassa* (Nc), *Sclerotinia sclerotiorum* (Ss), *Trichoderma reesei* (Tr) and *Ustilago maydis* (Um) as well as *Pyricularia* isolates collected from *O. sativa* (PoOs), *T. aestivum* (PoTa), *D. sanguinalis* (PoDs), *S. viridis* (PoSv), and *E. indica* (PoEi). Log2 Copy Number presents variation of copy number with increased red color means increased number of CAZymes.

**Additional files 5: Figure S2.** GH28 of P1609 (P1609_11576, P1609_5879, P1609_2497, P1609_5781, P1609_680 and P1609_5514)) and PoOs (MGG_09608, MGG_08752 and MGG_08938), PoDs (Ds0505_9820). Extra copies of GH28 in P1609 is marked by blue.

